# QRS Detection by Rules Based Multiple Channel Combinatorial Optimization

**DOI:** 10.1101/2023.06.16.545354

**Authors:** Bruce Hopenfeld

## Abstract

A multiple channel QRS detector is described. Separately for each channel, the detector generates sequences of peaks and statistically scores them according to: 1) peak prominence; 2) temporal regularity; 3) peak shape similarity; and 4) number of skipped beats. In the case of unstructured rhythms, the temporal regularity score is null and does not contribute to sequence quality. If at least one winning score from any channel exceeds a quality threshold, multi-channel sequences are generated from the winning sequences’ peaks and scored according to the above measures and peak time coherence across channels. The winning multi-channel sequence is then selected. The algorithm was applied to both channels of the MIT-BIH Arrhythmia Database. Over the entire database, the sensitivity (SE) and positive predictive value (PPV) were 99.93% and 99.96% respectively. For record 203, generally considered the most difficult one in the database, the SE and PPV were 99.80% and 99.76% respectively. The present algorithm fits within the framework of a previously described algorithm that can detect sinus rhythm in high noise conditions (e.g. waist based textile electrode recordings of a jogging subject).

## 1. Introduction

QRS detection based on combinatorial optimization has been applied to a number of datasets dominated by sinus rhythm [1-4]. In these works, an algorithm known as TEPS (Temporal Pattern Search), takes advantage of the relative temporal regularity of sinus rhythm to distinguish heartbeat sequences from noise. This work extends multiple channel combinatorial optimization to unstructured heart rhythms within an algorithmic framework that can successfully process very noisy signals. Such signals are likely to result in false positives when processed with a conventional serial QRS detector.

Such detectors are generally premised on the expectation that peak size/shape will very likely distinguish QRS complexes from noise. This expectation is satisfied in the context of the MIT-BIH Arrhythmia Database [5,6], which serves as a primary resource for testing QRS detection algorithms. Thus, relatively few false positives result when testing these algorithms on this database. However, in high noise conditions, QRS and noise peak shapes and sizes frequently overlap, which will result in false positives if these are the sole discrimination criteria. Adding a rhythm constraint, however, can enable detection of structured rhythms in such conditions [1-4].

The utilization of rhythm information is implicit in some neural networks [7] and explicit in recent publications from Nathan and Jafari [8], Li et al. [9], Modak et al.[10], and Alizadeh et al.[11] The former two approaches rely on formal algorithmic structures, particle filtering and Dynamic Bayesian Networks (DBN) respectively, whereas the present algorithm is rules based. Additional differences include: 1) TEPS’ use of a peak pair prominence ratio that acts as a local measure of signal to noise ratio (SNR) [2-4]; 2) TEPS’ tiered sequence search methodology; 3) TEPS’ hierarchical approach to obtaining a final set of QRS detections from segments; 4) TEPS’ RR Interval/Segment score clustering approach to select from amongst a potentially large number of leads (e.g. in the fetal ECG context).

Modak et al.[10] eliminate false peaks based upon RR interval criteria rather than basing peak detection itself on such criteria. Alizadeh et al.[11] describe a serial peak detection that estimates peak probability based in part on prior RR intervals.

In addition to the incorporation of rhythm information, the application of peak time coherence across channels can help to distinguish QRS complexes from noise [2, 4]. Yet, multiple channel QRS detectors are sparsely represented in the literature. Li et al. [9] integrated both channels of the MIT-BIH in a DBN, although they did not describe the methodology for identifying corresponding peaks across channels. TEPS explicitly incorporates peak time coherence information and on-the-fly generation of inter-channel peak time offsets [2,4]. In addition to the advantage of peak time coherence information, multiple channel implementations can allow one channel to compensate for another that is suffering from temporary bursts of noise or signal dropout.

Convolutional neural networks (CNNs) have achieved the best single channel results on the MIT-BIH database [7, 9]. However, it is not clear whether CNNs can generalize to high noise conditions [12]. In addition, CNNs tend to have relatively poor temporal resolution.

The advances of the present algorithm include

- accurate detection of QRS complexes in low/medium noise conditions, in which conventional QRS detection algorithms operate, while having the capability to seamlessly provide high temporal resolution sinus rhythm QRS detection in high noise conditions;
- implementation of peak time coherence based combinatorial optimization across multiple channels for both structured (e.g. sinus) and unstructured (e.g. atrial fibrillation) heart rhythms, and for abnormal QRS shapes;
- a tiered search and quality estimation strategy that ensures that low noise segments are efficiently processed;
- use of rhythm structure across entire 5-10 second long segments, so that, for example, segments that include multiple PVCs are more easily distinguished from noise.

### 2. Algorithm Overview

Figure 1 is a high-level block diagram of the algorithm, which processes data in overlapping 8 s segments that are adaptively extended to 10 s in the case of bradycardia. For convenience, the algorithm will be described with reference to two channels, but it may be extended to any number of channels. The first five steps, outlined by a dashed line, are performed separately for each channel. For each segment, pre-processing (filtering etc.) is performed. In this step, a noise analysis is performed by computing a discrete cosine transform and checking the autocorrelation of a second derivative signal. Very noisy segments may contain some useful information and are not rejected, but signals characterized by a power spectrum far outside the normal for humans are excluded. Also, relatively large periodic noise in a frequency range beyond the human maximum heart rate must be detected and segment quality adjusted accordingly (because such noise can produce false positives in the context of the present algorithm’s structured rhythm sequence search methodology [1,2]).

**Figure 1.**
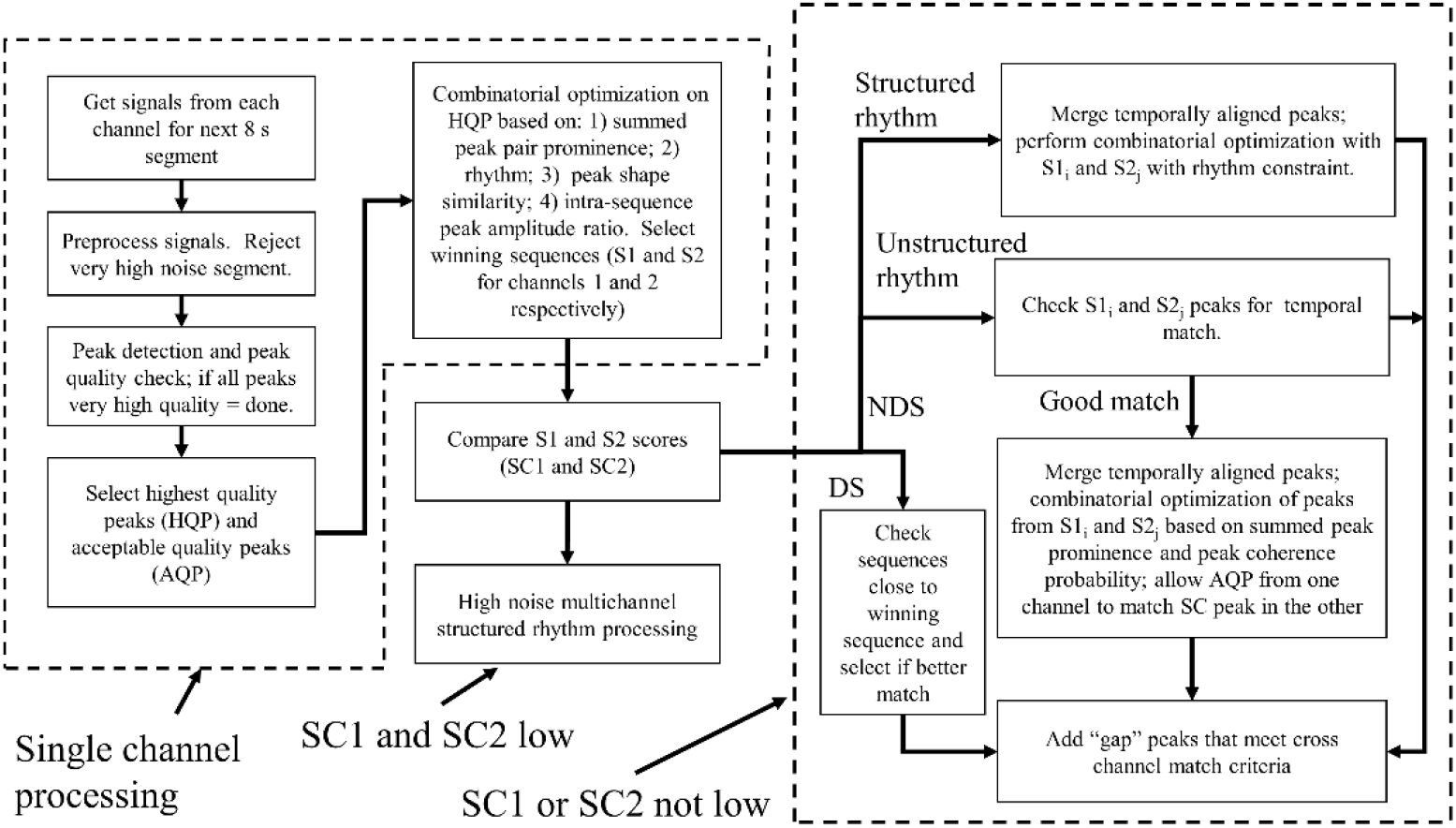
High level multiple channel flowchart. NDS represents the non-dominant sequence branch; NS represents the dominant sequence branch.

In the next step, the algorithm detects peaks. If the peaks are of uniformly high quality based on peak prominence (Section 2.6) and segment based peak amplitude ratios, they are selected as a final sequence and the processing for the current segment is complete. Otherwise, it selects two sets of peaks, relatively high quality peaks (HQP) and a disjoint set of acceptable quality peaks (AQP). If there is no clear delineation of the highest quality peaks, then the N (e.g. 17) best peaks are selected as HQP. The number of AQP is limited to a fixed amount (e.g. 13). In the next step, the algorithm performs combinatorial optimization on the HQP with an objective function based on: 1) summed peak pair prominence [2]; 2) rhythm (Section 2.2); 3) peak shape similarity [14]; and 4) intra-sequence peak amplitude ratio (minimum amplitude/median amplitude). The best sequences are selected (S1 and S2 for channels 1 and 2 respectively) that have associated scores S1 and S2 respectively.

If SC1 and SC2 are both low, indicative of low signal to noise in both channels, then processing branches to the high noise multichannel sinus rhythm methodology described in previous work [2,4]. Otherwise, if at least one of SC1 or SC2 is at least medium quality, there are three processing branches.

If the quality of one of the sequences is much higher than the quality of the other, the dominant sequence branch (DS in the figure) is taken. The dominant channel’s sequences are checked to determine whether there are sequences with a score close to that of the dominant sequence and that have the same length. If so, it is possible that one of these sequences is higher quality than the winning sequence when peak matching with the other channel’s best sequence is considered. Peak matching coherence scores are added to both the winning sequence and the close sequences, and a new highest scoring sequence is selected. In the next step, this winning sequence is checked for “gaps”, which will be described more fully below.

Returning to the last branch point, if the scores SC1 and SC2 are reasonably close, the non-dominant sequence branch (NDS in the figure) is taken, which leads to another branch point. The algorithm checks whether the rhythm associated with S1/S2 is structured or not. If the rhythm is structured, the structured rhythm branch is taken, and peaks that are temporally aligned across channels are merged as described in [2]. The AQP from one channel are allowed to match an HQP peak in the other channel, thereby increasing the score of any sequence that includes that peak. For AQP peaks, the peak coherence score is weighted by AQP amplitude. Combinatorial optimization is performed, and sequences are assessed for quality according to measures that include rhythm structure (Section 2.2).

Returning to the rhythm branch point, if structured rhythm is not indicated, then the sequence’s peaks are checked for timing coherence [2, 4]. If very few peaks temporally align/match, one channel’s sequence is likely of poor quality, and the highest scoring sequence is selected and checked for gaps. Otherwise, if a sufficient number of peaks match, then sequences are generated that include all of the matched peaks (core peaks) and combinations of a subset of the remaining (non-matching) peaks from both S1 and S2. For a remaining peak to qualify for this subset, it must be: 1) of high quality (Section 2.6) (above that required for classification as an HQP), and/or 2) matched with an AQP from the other channel. The resulting sequences are scored according to 1) summed peak pair prominence (summing across channels [2, 4]); and 2) peak coherence probability [2, 4]. The winning sequence is selected and checked for gaps.

The gap filling step is again performed in case S1 and S2 do not include all high quality peaks that fit into high quality sequences. In the case of structured rhythm, “gaps” can be filled only by high quality interpolated PVC peaks that temporally align across channels. In the case of unstructured rhythm, gap filling is a backstop measure that examines HQP in both channels that were not included in S1/S2 but that nonetheless fit into S1/S2 and qualify as a QRS when peak time coherence is included in the assessment of peak quality. (S1 and S2 are single channel sequences whose scores were not based on peak timing coherence.) A final step, which was not included in the present algorithm but may be necessary in a broader dataset, would be to check the score of the final winning sequence to ensure that it is of sufficiently high quality. Otherwise, the high noise structured rhythm block would be invoked.

Figure 2 is a flow chart of single channel processing. After preprocessing, a noise analysis is performed by checking the peak prominence of the largest peaks (Section 2.6). If the peak prominence scores indicate a high SNR condition, then the high quality peaks are examined to determine whether they are associated with P/T waves. If a threshold number of peaks cluster tightly within time ranges, relative to larger peaks, that are consistent with P/T wave timing, the peaks within the clusters are removed from further processing. Next, from either the first block or the P/T block, HQP are checked for likelihood of a structured rhythm and an associated average segment RR interval RRP. (In the case of e.g. a bigeminal rhythm, there may be one RR interval associated with the segment.) This is performed by autocorrelation of a peak space signal generated from the HQP [13].

**Figure 2.**
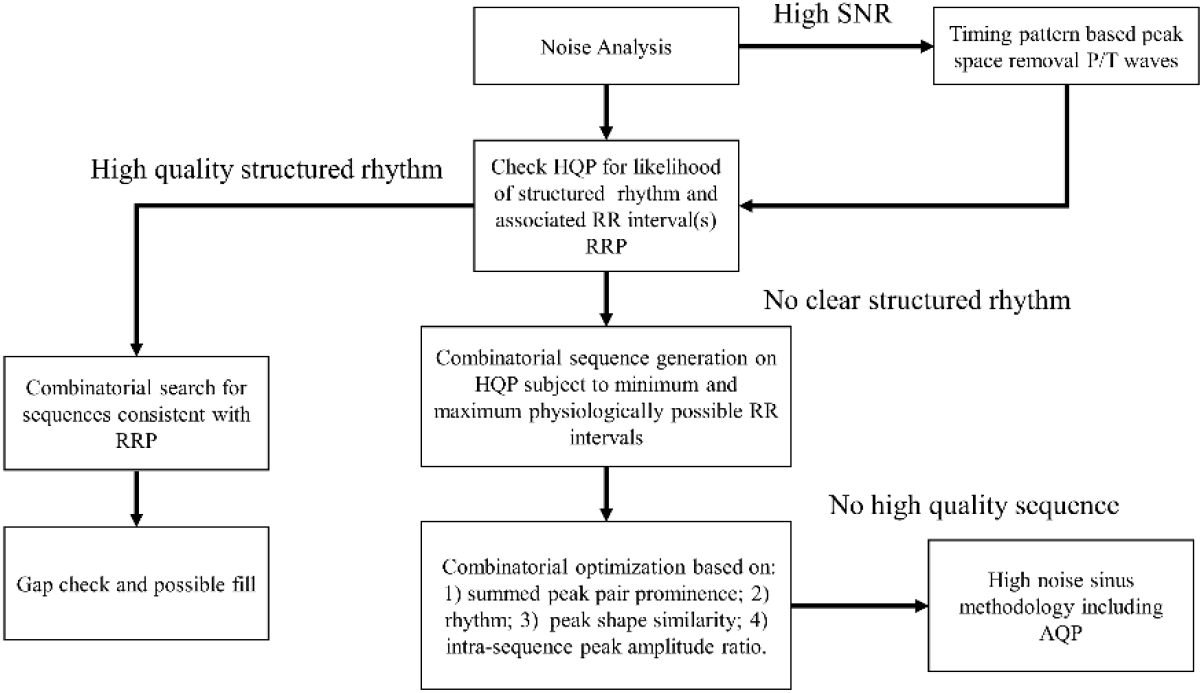
High level single channel flowchart.

If a structured rhythm is indicated, then a combinatorial search is performed such that the generated sequences are consistent with RRP. If there are any gaps in the best sequence, they are examined to determine whether to fill them in with an AQP. Gaps may be filled in with sinus rhythm beats or PVCs. Intra-sinus beat gaps are checked for the possibility of interpolated PVCs, whose peaks are selected only if they are relatively high quality.

Returning to the structured rhythm check, if structured rhythm is not apparent, then an unconstrained sequence search is performed, and sequences are scored according to previously mentioned criteria. These criteria include a rhythm score, which is set to null in the case of sequences with an unstructured rhythm. (The segment may have a structured rhythm sequence that was not detected by the previously described high level structured rhythm check.) If there are no high scoring sequences, then high noise sinus rhythm methodology is invoked. This step is shown in the single channel processing associated with Figure 2 but it may occur after the checking of scores SC1 and SC2 in the multi-channel processing (Figure 1).

### 2.1 Peak Pair Prominence

Compared to noise peaks, pairs of heart beats are more likely to be separated from one another by “smaller” peaks. A Bayesian analysis of peak pair prominence is detailed in [2], which also shows the summation of peak pair prominence scores across channels. In the present work, the peak pair prominence measure has been modified for the purposes of multiple channel scores, so that it is an increasing function of RR interval if a single channel sequence (S1/S2) has a sufficiently high score. (Herein, inter-peak intervals will be referred to as RR intervals even for peaks that are not QRS complexes.) In particular, if peaks are closer together (lower RR interval), they are relatively less likely to be interrupted by other peaks compared to peaks separated by a longer RR interval; thus, a smaller RR interval carries a lower evidentiary value from the standpoint of peak pair prominence. Increasing the peak pair prominence measure (score) as a function of increasing RR interval reflects this prior information. (It is also possible that signal dropout from, e.g., bad electrode contact, can cause long stretches of seemingly noise free signal. This can possibly be handled by checking impedance/resistance. However, even absent such external information, on balance, the above assumption is justified, at least in context of the MIT-BIH database.)

An example of peak pair prominence summing will be described with reference to Figure 3. In channel 1, two possible subsequences including peaks a and c are {a,b,c} and {a,e}. In channel 2, a single subsequence involving peaks d and e is simply {d,e}. Peaks a and d temporally align and are thus merged. Similarly, peaks c and e are merged. The summed peak prominence score (PPS) for multichannel sequence {a/d, c/e} is simply the sum of the single channel peak pair ratios (PPR) across channels 1 and 2: PPR(a,c) + PPR(d,e). The summed peak prominence score (PPS) for multichannel sequence {a/d, b, c/e} is simply the sum of the single channel PPR’s from channel 1: PPR(a,b) + PPR(b,c). (Because peak b has no match in signal 2, there are no corresponding peak pairs in channel 2.) The PPS for {a/d, c/e} will be greater than the PPS for {a/d, b, c/e} if PPR(a,c) + PPR(d,e) is greater than PPR(a,b) + PPR(b,c). This is indeed the case, because of the short RR interval between peak b and c which in turn reduced the raw PPR(b,c), and peak b was properly rejected by the algorithm. Conversely, the peak in channel 2 labelled as “QRS” in the figure was properly accepted.

**Figure 3.**
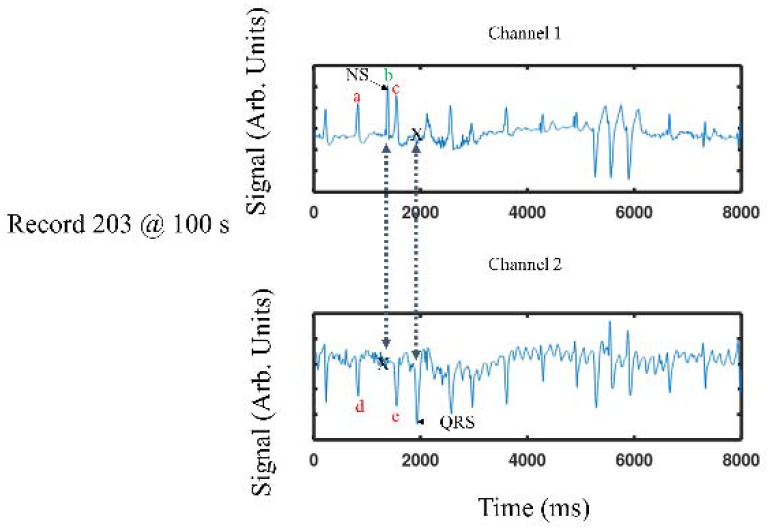
Examples of the computation of peak pair prominence ratio across two channels. In channel 1, peak b is a noise spike and there is no corresponding peak in channel 2, as indicated by the arrowed dashed line and the “X” in channel 2. Conversely, channel 2 has a true QRS, labelled as such, that does not have a corresponding peak/QRS in channel 1, as indicated by the arrowed dashed line and the “X” in channel 1. The peak pair prominence ratios of subsequence {a,c} and {d,e} are added. See text for details.

The minimum peak pair prominence ratio across all peak pairs in a sequence is a useful metric for assessing whether a sequence satisfies stopping criteria so that no further processing is necessary.

### 2.2. Rhythm

The temporal regularity of certain heart rhythms can help distinguish true heart beat sequences from sequences of noise. References [1-4] involve a rhythm score that is based on the summed change in successive RR intervals in a sequence, taking into account skipped beats. This score fails to distinguish higher order rhythm patterns such as multiple PVCs separated by sinus beats or bigeminal rhythms. To account for these types of rhythms, a search is performed for a short-long beat pattern with short beat RR intervals that cluster within a certain range. For qualifying sequences, the short beats are effectively treated as skips for the purposes of the temporal regularity calculation [1,2] but do not count against the sequences as true skips.

To account for sequences with a mix of structured and unstructured rhythms, changes in RR intervals that exceed a threshold are removed from the temporal regularity calculation and treated as a type of skip that is linearly penalized. If the overall temporal regularity score falls below a specified threshold, then it is set to 0; below this threshold, the temporal regularity measure adds no predictive value, so that degrees of irregularity underneath this threshold are effectively ignored.

The above approach worked well for the MIT-BIH database, but for future work, a more general approach, such as one of the following, may prove beneficial across a larger sample of rhythm types: 1) clustering analysis on the Poincare plots for different sequences; 2) neural network assessment of the RR interval patterns.

### 2.3 Peak Shape

Intra-sequence QRS complex shape similarly can help to distinguish true peaks from noise [14] and is part of a sequence’s score. Multipoint first derivative signals are compared to one another through a cross correlation aligned by primary peak time.

### 2.4 Peak Time Coherence

Heart beats will occur close in time across channels whereas noise peaks (in uncorrelated channels) will tend to temporally align across channels only by chance. Figure 4 shows an example of peak time coherence increasing the likelihood that a peak is a QRS complex. A Bayesian analysis of peak time coherence is described in [2]. In the MIT-BIH, the irregular shapes of many types of QRS complexes result in a more challenging temporal alignment problem than sinus rhythm beats. To account for this, peak times were modified for certain types of closely spaced peaks, as will be described in Section 2.5.

**Figure 4.**
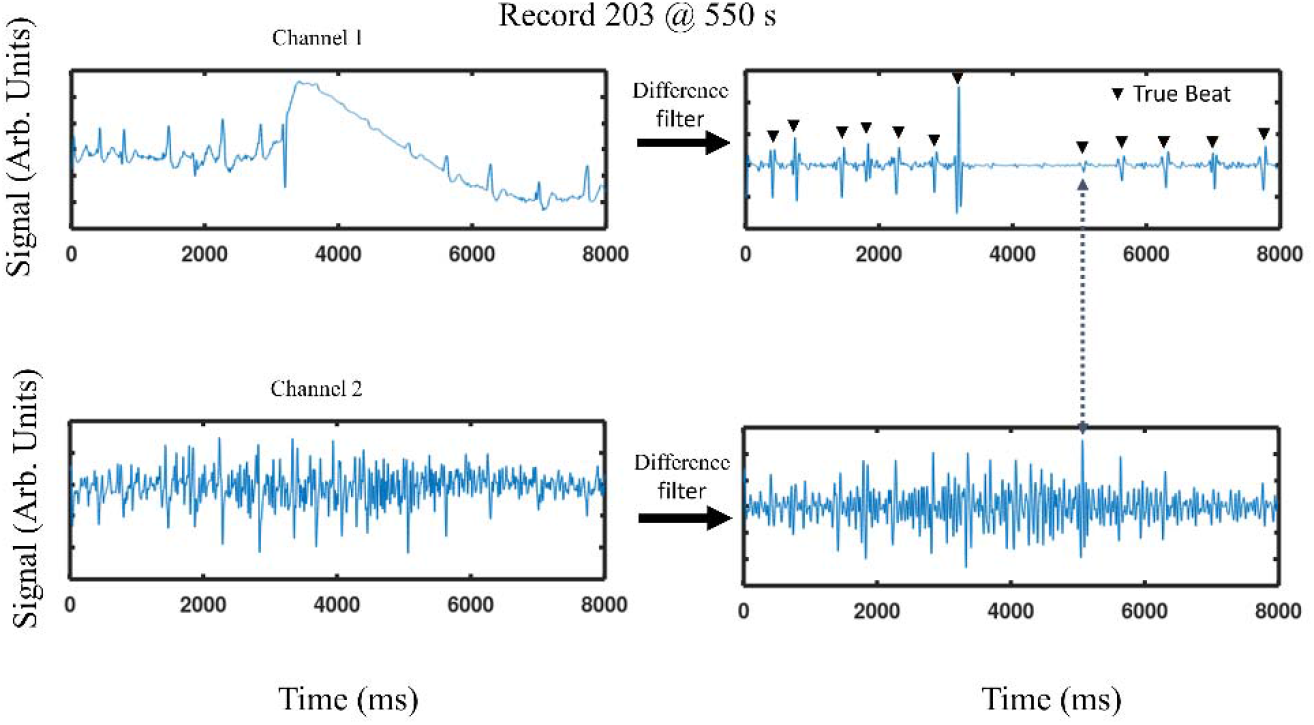
An example of a noisy segment (channel 2) that nonetheless carries useful information that is used to properly detect an ambiguous QRS complex indicated by the arrowed dashed line. Note that in this case, for channel 2, the multi-point second derivative may actually enhance noise. “True beat” means a peak annotated as such in the MIT-BIH database. (Three peaks in this segment in the gap between approximately 3500ms and 5000ms, not annotated as true beats, were nearly detected as beats by the present algorithm.)

Also, in the low/medium noise processing blocks, cross channel peaks were matched with two different peak time offsets, whichever produced the best match: 1) an overall majority offset, determined as in previous work; and 2) an automatically assumed 0 offset. Instead of match quality being a decreasing function of a rather short time window (e.g. 10-20ms), a uniform quality was assigned within a larger match window of 80 ms. Further, in the gap filling stage, if a winning sequence is missing a peak that corresponds to peaks in both S1 and S2 within a larger (e.g. 150 ms) match window, the S1/S2 peaks are considered a match, merged, and added to the winning sequence.

### 2.5 QRS Isolation/Merging Peaks

In low noise conditions, the separation of QRS complexes from the surrounding signal is straightforward, even when a QRS is fractionated. In high noise conditions, however, it is more difficult to tag a group of peaks as a particular QRS complex, so to attain high temporal resolution for sinus rhythm RR intervals, it is often necessary to associate closely spaced peaks with different (potential) QRS complexes. However, if this separation is applied to a group of peaks that in fact corresponds to a single QRS complex, it will confound the peak quality/peak prominence scores because a single QRS complex will appear as noise to itself. Further, fractionated, slurred QRS complexes can confound peak time coherence measures.

To handle the full range of possible noise conditions, the decision whether to merge closely spaced (within physiological QRS duration) peaks is a function of the noise level in the time interval close to the peaks. If the subject group of peaks has prominence over the peaks just before and after it, and the group of peaks have similar amplitudes, then they are merged. The time of the merged peak is taken as the midpoint of a summed/averaged measure of the absolute value of the second derivative over the period encompassing the group of peaks, akin to many serial QRS detectors. This type of merger is not performed if one peak has a substantially greater amplitude than the others, in which case that dominant peak’s time is taken as the peak time for the group. However, the other peaks in the group are not counted as noise peaks against the primary peak for the purposes of peak prominence.

### 2.6 Peak Quality Based on Individual Peak Prominence/Amplitude Envelope Adjustment

Segment quality is assessed, in part, with regard to the quality of individual peaks, which in turn is a function of their peak prominence. Individual peak prominence for a peak *j* at time *t*_*j*_ is:

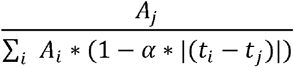

where *A*_*i*_ is the amplitude (in multipoint second derivative space) of the ith peak, α is a parameter that controls the decay of the influence of adjacent peak amplitude, and the sum is taken over all peaks that have a non-zero influence over peak *j*. The parameter α can be adjusted to attain a relatively more global or local measure of peak prominence.

In some segments, for example the one shown in the upper panel in Figure 4, the signal amplitude changes due to an impedance change, imparting a varying envelope on the signal. It is desirable to correct peak amplitudes according to the envelope. A difficulty arises in distinguishing an impedance change from cardiac activity; for example, lower amplitude noise spikes or T waves in the context of low heart rates, especially bradycardia, can appear as a changing envelope amplitude. In the absence of an exogenous measure of impedance, the envelope amplitude adjustment was made only for the purposes of the peak space structured rhythm check. (In some instances, features of the signal may enable the envelope to be extracted even in the case of unstructured rhythm, e.g. in the case where there are closely spaced similarly shaped QRS complexes, but this methodology was not implemented.)

### 2.7 Merging Winning Segment Sequences

Segments are typically 8 seconds long with a 3 second overlap. To generate a final set of detected peaks, the winning sequences from each segment must be merged. If the overlapping detections match in time, the corresponding peaks are selected. If there is a conflict between overlapping peaks, then the peaks that result in the most temporally regular overall sequence are selected.

### 2.8 Processing Details

Raw signals are preprocessed by low pass filtering with a 5th order Butterworth filter with a cutoff frequency of 45Hz. The filtered signals are downsampled to 256Hz, and the resulting signal x() is differenced according to: y(i) = x(i-12)-2*x(i-6)+x(i). A secondary check is made to ensure that, in the case of a large second difference, there is a first derivative polarity change of sufficient magnitude to qualify as a peak (to handle the case of a sharp change in baseline). All computations were performed on a 2017 Hewlett Packard Laptop with an Intel Core i3-8130U CPU, base frequency 2.20GHz, with 8GB of RAM.

## 3. Databases; Parameter Choice

The algorithm was run on both channels of the MIT-BIH Arrhythmia Database. The algorithm was refined, and various parameters were heuristically chosen, by examining various segments across a variety of records within the database. The parameters were not optimized.

## 4. Results

Over all records in the MIT-BIH database, sensitivity (SE) and positive predictive value (PPV) were 99.93% and 99.96% respectively. SE and PPV for individual records is set forth in Table 1.

**Table 1.**
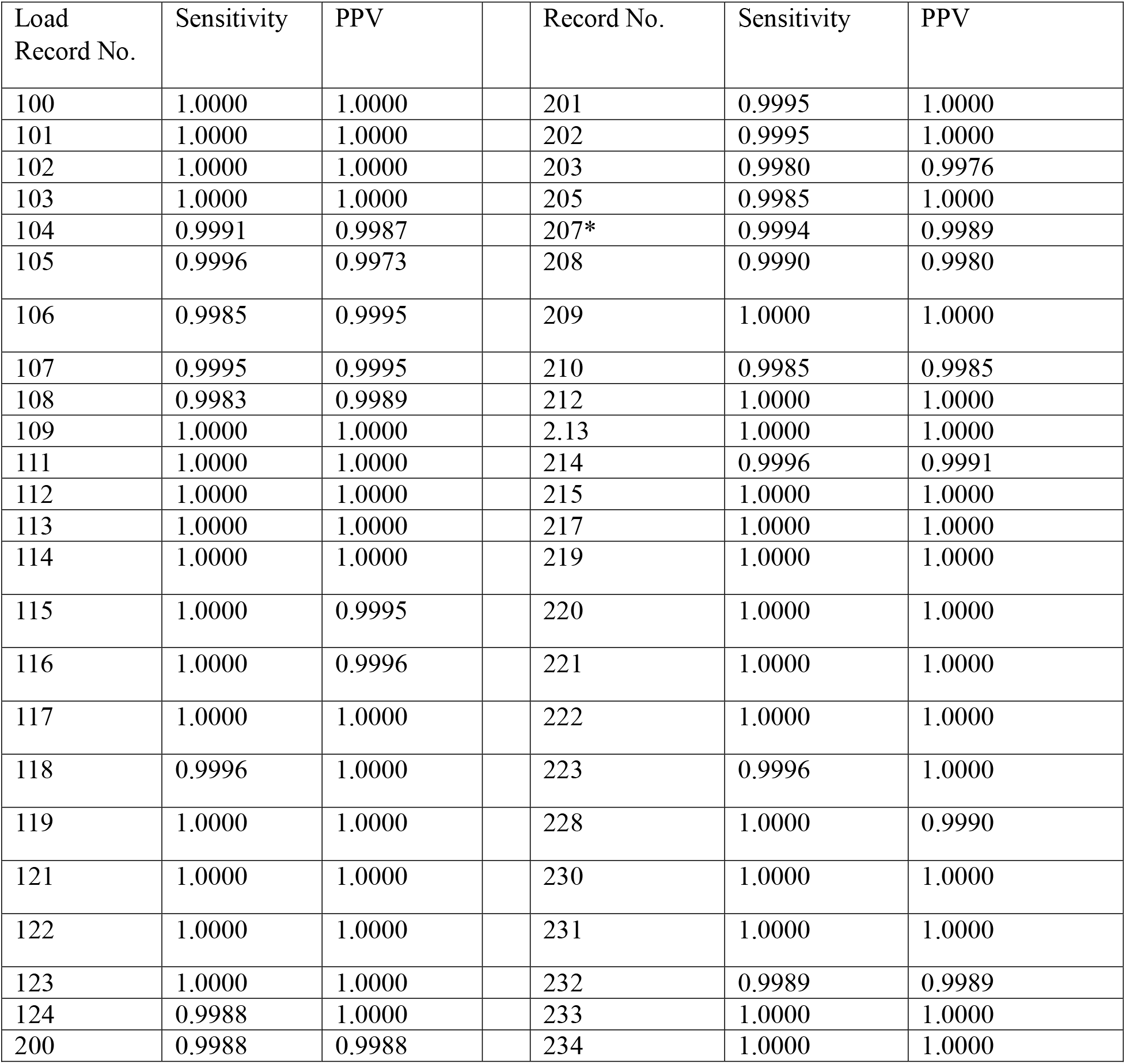
MITBIH Results By Record* For record 207, any 5 second sub-segment that includes ventricular flutter was excluded.

| Load Record No. | Sensitivity | PPV |  | Record No. | Sensitivity | PPV |
| --- | --- | --- | --- | --- | --- | --- |
| 100 | 1.0000 | 1.0000 |  | 201 | 0.9995 | 1.0000 |
| 101 | 1.0000 | 1.0000 |  | 202 | 0.9995 | 1.0000 |
| 102 | 1.0000 | 1.0000 |  | 203 | 0.9980 | 0.9976 |
| 103 | 1.0000 | 1.0000 |  | 205 | 0.9985 | 1.0000 |
| 104 | 0.9991 | 0.9987 |  | 207* | 0.9994 | 0.9989 |
| 105 | 0.9996 | 0.9973 |  | 208 | 0.9990 | 0.9980 |
| 106 | 0.9985 | 0.9995 |  | 209 | 1.0000 | 1.0000 |
| 107 | 0.9995 | 0.9995 |  | 210 | 0.9985 | 0.9985 |
| 108 | 0.9983 | 0.9989 |  | 212 | 1.0000 | 1.0000 |
| 109 | 1.0000 | 1.0000 |  | 213 | 1.0000 | 1.0000 |
| 111 | 1.0000 | 1.0000 |  | 214 | 0.9996 | 0.9991 |
| 112 | 1.0000 | 1.0000 |  | 215 | 1.0000 | 1.0000 |
| 113 | 1.0000 | 1.0000 |  | 217 | 1.0000 | 1.0000 |
| 114 | 1.0000 | 1.0000 |  | 219 | 1.0000 | 1.0000 |
| 115 | 1.0000 | 0.9995 |  | 220 | 1.0000 | 1.0000 |
| 116 | 1.0000 | 0.9996 |  | 221 | 1.0000 | 1.0000 |
| 117 | 1.0000 | 1.0000 |  | 222 | 1.0000 | 1.0000 |
| 118 | 0.9996 | 1.0000 |  | 223 | 0.9996 | 1.0000 |
| 119 | 1.0000 | 1.0000 |  | 228 | 1.0000 | 0.9990 |
| 121 | 1.0000 | 1.0000 |  | 230 | 1.0000 | 1.0000 |
| 122 | 1.0000 | 1.0000 |  | 231 | 1.0000 | 1.0000 |
| 123 | 1.0000 | 1.0000 |  | 232 | 0.9989 | 0.9989 |
| 124 | 0.9988 | 1.0000 |  | 233 | 1.0000 | 1.0000 |
| 200 | 0.9988 | 0.9988 |  | 234 | 1.0000 | 1.0000 |

## 5. Discussion

There are very few multiple channel QRS detectors with which to compare the present results. Li et al. [9] report a double channel SE and PPV of 99.78% and 99.74% for the MIT-BIH database. A neural network has achieved an SE/PPV of 99.94% and 99.97% [7] on a single channel of the MIT-BIH database, slightly bettering the double channel results in the present study. As described in the Introduction, neural networks have various limitations.

The present algorithm detects whether a segment, across all channels, has a noise level that is too high to accurately detect unstructured rhythms. Thus, these high noise segments will likely not produce false positive detections. Instead, either a sinus (structured) rhythm is found, or the segments are discarded as noise (or possibly analyzed via histograms for multi-segment statistics). This selectivity is likely not possible with any type of serial QRS detector, which dominate the non-neural network approaches to QRS detection.

It is not clear how convolutional neural networks would handle high noise conditions, which likely require some sort of combinatorial optimization. In such conditions, there is a large overlap in the shape/size of QRS complexes, and there are frequently “orthogonal” segment sequences that have similar likelihoods, in which case it is necessary to separate them and track the associated RR intervals over longer periods. Convolutional neural networks do not lend themselves to combinatorial optimization, which has been the province of graph neural networks. Also, the CNN approach, as generally applied to date, would appear to have relatively poor temporal resolution. Finally, the generalizability of CNNs remains an open question.

## Notes

### Competing Interest Statement

The authors have declared no competing interest.

